# Sirt5 deficiency causes post-translational protein malonylation and dysregulated cellular metabolism in chondrocytes under obesity conditions

**DOI:** 10.1101/2020.11.30.404103

**Authors:** Shouan Zhu, Albert Batushansky, Anita Jopkiewicz, Dawid Makosa, Kenneth M. Humphries, Holly Van Remmen, Timothy M. Griffin

## Abstract

**Objective:** Obesity accelerates the development of osteoarthritis (OA) during aging and is associated with altered chondrocyte cellular metabolism. The objective of this study was to investigate the role of sirtuin 5 (SIRT5) in regulating chondrocyte protein lysine malonylation (MaK) and cellular metabolism under obesity-related conditions.

**Methods:** MaK and SIRT5 were immunostained in knee articular cartilage of obese *db/db* mice and different aged C57BL6 mice with or without destabilization of the medial meniscus (DMM) surgery to induce OA. Primary chondrocytes were isolated from 7-day-old WT and *Sirt5*^*−/−*^ mice and treated with varying concentrations of glucose and insulin to mimic obesity. Sirt5-dependent effects on MaK and metabolism were evaluated by Western blot, Seahorse Respirometry, and gas/chromatography-mass/spectrometry (GC-MS) metabolic profiling.

**Results:** MaK was significantly increased in cartilage of *db/db* mice and in chondrocytes treated with high concentrations of glucose and insulin (Glu^hi^Ins^hi^). Sirt5 protein was increased in an age-dependent manner following joint injury, and Sirt5 deficient primary chondrocytes had increased MaK, decreased glycolysis rate, and reduced basal mitochondrial respiration. GC-MS identified 41 metabolites. Sirt5 deficiency altered 13 distinct metabolites under basal conditions and 18 metabolites under Glu^hi^Ins^hi^ treatment. Pathway analysis identified a wide range of Sirt5-dependent altered metabolic pathways that include amino acid metabolism, TCA cycle, and glycolysis.

**Conclusion:** This study provides the first evidence that Sirt5 broadly regulates chondrocyte metabolism. We observed changes in Sirt5 and MaK levels in cartilage with obesity and joint injury, suggesting that the Sirt5-MaK pathway may contribute to altered chondrocyte metabolism that occurs during OA development.

## INTRODUCTION

Osteoarthritis (OA) is a leading cause of chronic pain and disability worldwide, affecting 9.6% of men and 18.0% of women aged ≥ 60 years (1). Aging is the primary risk factor for OA; however, increased longevity alone is not sufficient to explain the escalating prevalence of OA in our population. A recent study of long-term trends in OA prevalence in the United States strongly suggests that factors other than increased longevity are responsible for a doubling in the prevalence of OA since the mid-20^th^ century (2). One potential contributing factor is the increased prevalence of type 2 diabetes and other metabolic conditions contributing to metabolic syndrome (MetS) (3, 4). Although mechanical stress on joints is a contributing factor to obesity-associated OA, obesity and MetS increase the risk of hand OA (5, 6), indicating that systemic metabolic factors also impair joint health.

Systemic metabolic diseases such as type 2 diabetes adversely affect various cellular processes, such as mitochondrial bioenergetics, nutrient sensing, glycolysis, and proteostasis (7). Consistent with these impaired cellular processes, there is broad evidence that impaired cellular metabolism and mitochondrial dysfunction is a general feature of OA chondrocytes (8, 9). For example, impaired electron transport chain activity in OA chondrocytes increases cartilage oxidative stress (8). In addition, reduced activity of cellular energy sensors (e.g., AMPK and sirtuins) is also implicated in OA pathogenesis (10–12). Despite these important findings, there is a critical knowledge gap in understanding how systemic metabolic conditions, such as type 2 diabetes, impair joint tissue cellular metabolism.

One theory linking systemic MetS-related conditions to altered cellular function is the induction of ‘cellular carbon stress’ (13, 14). Excessive cellular glucose and lipid uptake causes a cellular carbon overload that leads to the accumulation of numerous reactive acyl-group metabolites (14). These metabolites post-translationally modify hundreds of proteins (PTMs), altering their enzymatic activity (13). Many forms of acyl-group PTMs exist, including lysine malonylation, succinylation, and acetylation (13). These modifications have been shown to impair cellular function during the development of obesity and aging-associated diseases. For example, increasing the acetylation of proteins in the liver that regulate fatty acid oxidation accelerates the development of MetS (15). The role of these PTMs in cartilage metabolism and oxidant defense remains largely unknown. We previously reported that an age-related increase in the post-translational lysine acetylation of the mitochondrial antioxidant SOD2 increased the susceptibility of cartilage to oxidative stress (11). Importantly, these changes were associated with a reduction in Sirtuin 3 (11), the primary NAD^+^-dependent lysine deacetylase for mitochondrial proteins.

Mammals possess seven sirtuins (Sirt1-7) that modulate distinct metabolic and stress response pathways by their deacylase activity. Sirt5 was initially reported as a lysine deacetylase found in both mitochondria and cytoplasm. However, recent studies have shown it has low deacetylase activity and strong demalonylase and desuccinylase activity (16). Label-free mass spectrometry of immunoprecipitated malonyl-lysine peptides showed that Sirt5 deficiency enriched the hypermalonylation (MaK) of liver proteins regulating glycolysis (17). Consistently, *Sirt5*^*−/−*^ hepatocytes showed decreased production of lactic acid and reduced glycolytic flux (17). The biological consequence of protein malonylation has not been extensively studied; however, recent studies strongly suggest that enhanced lysine malonylation contributes to metabolic disturbances associated with diabetes. For example, a large-scale analysis of protein malonylation in the livers of *db/db* mice, a type 2 diabetes mouse model, showed upregulated lysine malonylation of metabolic enzymes involved in glucose catabolism and fatty acid oxidation (18).

The goal of the current study was to investigate the effect of Sirt5-mediated protein malonylation (MaK) on chondrocyte cellular metabolism and to evaluate the potential role of the Sirt5-MaK pathway in OA development. Chondrocytes primarily rely on glycolysis to provide cellular ATP. Thus, we hypothesized that Sirt5-mediated MaK regulates chondrocyte cellular metabolism under basal and obesity-associated conditions. To test this hypothesis, we first examined the level of MaK accumulation *in vivo* in the knee articular cartilage of obese mice or *in vitro* in primary chondrocytes treated with high glucose and high insulin culture conditions. We also examined the effect of aging and OA on cartilage Sirt5 expression. We then tested the functional effect of Sirt5 on chondrocyte metabolism using Seahorse Respirometry to measure the rates of glycolysis and oxidative phosphorylation. To better understand the broader effects of Sirt5 on the chondrocyte metabolic network, we used Gas/Chromatography-Mass/Spectrometry (GC-MS) to conduct a semi-targeted cellular metabolomics analysis of the relative changes in metabolites in wild-type (WT) and Sirt5-deficient primary chondrocytes. Thus, with multiple *in vivo* and *in vitro* model systems, we provide the first evidence for a role of Sirt5 in regulating chondrocyte metabolism and MaK under obesity and OA conditions.

## MATERIALS AND METHODS

### Animals and Experimental Design

All experiments were conducted in accordance with protocols approved by the AAALAC-accredited Institutional Animal Care and Use Committee at the Oklahoma Medical Research Foundation (OMRF). Male C57BL/6J WT (n = 4) and C57BL/6J background mice homozygous for the diabetes spontaneous mutation in the leptin receptor (*db/db*) (n = 4) were purchased from The Jackson Laboratory (Bar Harbor, ME, USA) and were acclimated within the OMRF vivarium for one week prior to conducting experimental procedures. Additional male mice (C57BL/6J background) were bred for evaluating the effect of age on injury-induced post-traumatic OA using the destabilization of medial meniscus (DMM) model as previously described (19). Surgeries were initiated at 14 weeks of age (n = 8) and 48 weeks of age (n = 8), and joint tissues were collected 8 weeks after surgery such that the final ages were 22 and 56 weeks. Mice were anesthetized during the surgical procedure using isoflurane, and post-surgical analgesia was provided by subcutaneous injection of buprenorphine (0.09 mg/kg). The contra-lateral limb was evaluated as a non-surgical control. Male and female Sirt5 heterozygous knockout mice (B6;129-Sirt5^tm1FWa^/J, Stock No: 012757) were purchased from The Jackson Laboratory after Cryo Recovery of embryos (20). The mice were then bred to generate both WT and Sirt5 homozygous knockout (*Sirt5*^*−/−*^) mice. All mice were group housed (≤5 animals/cage) in ventilated cages in a temperature-controlled room maintained at 22 ± 3°C on 14h:10h light/dark cycles with *ad libitum* access to chow and water.

### Immunohistochemical staining for MaK or Sirt5 on mouse knee joint sections

Knee joint sections utilized for MaK and SIRT5 immunostaining were obtained as convenience samples from prior lab studies. Sagittal sections from age-matched 24-wk-old lean WT (n=4) and obese *db/db* mice (n=4) were used for MaK immunohistochemical staining. Isolated knee joints were fixed and embedded in paraffin as previously described, excluding micro-computed tomography analysis (21). Sections from 22- and 56-week-old mice that underwent DMM surgery were used for SIRT5 immunostaining. Isolated knee joints were fixed in freshly prepared 4% paraformaldehyde for 48 hours at 3°C, decalcified in Cal-Ex Decalcifier (Fisher Chemical) for 3 days at 3°C, dehydrated, and embedded in paraffin for coronal sectioning. Joint sections were deparaffinized, rehydrated, and incubated with antigen retrieval R-Buffer A (EMS, 62706-10) at 80°C for 1 hour. Slides were then treated with 2% H_2_O_2_, blocked using 5% BSA, and incubated overnight at 4°C with antibody anti-SIRT5 (Lifespan Biosciences, 1:100 dilution) or MaK (CST, #14942s, 1:200 dilution) overnight. Negative controls were processed in parallel on the slide with rabbit IgG. Staining was detected using goat anti-mouse secondary antibody conjugated with HRP (Horseradish Peroxidase) (Cell Signaling Technology, 7074, 1:250) with HIGHDEF Red IHC Chromagen HRP (Enzo, 11141807) for color development following manufacturer’s protocol. Sections were also counter-stained with hematoxylin for 30 seconds. Overall staining intensity for MaK and SIRT5 in uncalcified cartilage was evaluated using a semi-quantitative scale of cell-staining intensity as follows: 0 for non-stained, 1 for weakly-stained, and 2 for strongly-stained. For MaK quantification, tibia and femur were scored and analyzed separately. For SIRT5 quantification, femoral and tibial scores were averaged and analyzed separately for the lateral and medial compartments due to the injury primarily causing OA pathology in the medial compartment.

### Primary mouse chondrocyte isolation

Primary murine chondrocytes were isolated from 7 day old male or female wild type (WT) or Sirt5 deficient (*Sirt5*^*−/−*^) mice according to a previously published protocol (22). Briefly, both tibial and femoral cartilage tissue were dissected from the knee joint. The cartilage was then incubated in 3 mg/mL Collagenase D (Roche) in DMEM solution for two 45 minute periods and transferred to 0.5 mg/mL Collagenase D in DMEM solution supplemented with 3% Liberase TL (Sigma) overnight. The tissue was then homogenized by pipetting up and down to release and suspend the cells, and the homogenate was filtered through a 40 μm strainer to remove large debris. Passage 0 cells were cultured and expanded in 6-well plates in complete DMEM media (Life Technology, 10567014) supplemented with 10% fetal bovine serum (FBS) and 1% penicillin/streptomycin (P/S) at 37 °C and 5% CO2. Chondrocytes reached confluency after 3-4 days in culture. Chondrocytes were then trypsinized, counted, and reseeded as passage 1 cells for specific experiments described in detail in the following sections.

### Screening for metabolic conditions that increase MaK

We screened for culture conditions that increase protein lysine malonylation (MaK) in chondrocytes. Passage 1 murine primary chondrocytes were treated with media supplemented with various combinations of glucose (2.5, 10, and 20 mM) and insulin (0, 10, and 20 nM) for 24 hours in glass bottom Greiner 96 well plates. Cells were fixed with 4% paraformaldehyde for 10 min, permeabilized with 0.2% Triton-X 100, blocked with 5% bovine serum albumin (BSA), and incubated overnight with primary antibody against MaK (CST, #14942s, 1:200 dilution) and COL2 (abcam, ab185430, 1:200). Cells were then washed with PBS (3 x 5 min) and incubated with anti-Rabbit Alexa Fluor 488 and anti-Mouse Alexa Fluor 546 secondary antibodies for 1 hour at room temperature. Cells were counter-stained with DAPI and preserved in PBS. Fluorescent images of cellular MaK and COL2 staining were obtained at 40-fold magnification using confocal microscopy (Zeiss LSM-710), and images were analyzed in ImageJ for staining intensity and cell number quantification. Total signal intensity was measured separately for MaK and COL2 staining and normalized to cell number as determined by DAPI counts. The experiment included 9 independent biological replicates (n=9 WT mice) for each treatment condition.

### Western blotting analysis for MaK and Sirt5

Total protein from confluent passage 1 *Sirt5*^*−/−*^ and WT chondrocytes cultured in 6 cm dishes were collected in RIPA buffer containing 20 mM nicotinamide (Sigma) and protease inhibitors cocktail (ThermoFisher). Proteins were subjected to electrophoresis using NuPAGE 4-12% Bis-Tris gels (Life Technology), transferred to nitrocellulose membranes, and blocked for 30 min with Odyssey TBS blocking buffer (LI-COR). The membranes were then incubated overnight with primary antibodies for anti-MaK (CST, #14942s) or SIRT5 (CST, #8779) diluted in TBS blocking buffer supplemented with 0.1% Tween-20. The membranes were then incubated with IRDye 800CW goat anti-rabbit secondary antibody (926-32211, 1:15,000 dilution). The membrane was imaged by Odyssey CLx Imaging System. In total, 7 independent biological samples from each genotype were evaluated and quantified.

### Seahorse-based analysis of chondrocyte cellular metabolism

Passage 1 primary chondrocytes from WT and Sirt5^*−/−*^ mice were seeded in 24-well Seahorse microplates (60,000 cells per well) one day before testing. 4 wells across the plate (A1, B4, C3, and D6) were maintained cell-free to serve as blanks for background subtraction. The cartridge was also prepared one day before the assay by hydrating it in Seahorse XF calibrant at 37 °C in a dry incubator. The day of the test the hydrated cartridge was loaded with inhibitor solutions into each port according to the manufacturer instructions and following preliminary optimization testing (Glycolytic Rate Assay: Port A (0.5 μM Rotenone/Antimycin-A), Port B (50 μM 2-Deoxy-d-glucose); Mito Stress Assay: Port A (2 μM Oligomycin), Port B (1 μM FCCP), Port C (0.5 μM Rotenone/Antimycin-A). Cells were washed with Seahorse XF DMEM medium (pH 7.4, Agilent 103575-100) 3 times and incubated in Seahorse XF DMEM medium in 37 °C for 30 mins. Cells were then incubated with Seahorse XF DMEM medium supplemented with assay substrate (10 mM glucose for Glycolytic Rate Assay, 1 mM pyruvate for Mito Stress Assay). Tests were then conducted on a Seahorse XFe24 Analyzer (Agilent) following manufacturer instructions for the ‘Glycolytic Rate Assay’ or ‘Mitochondrial Stress Assay’ protocol. After test completion, the plates were collected to measure total protein concentration in each well using Pierce BCA Protein Assay Kit (#23225). Data were normalized to total protein content for comparison between WT and *Sirt5*^*−/−*^ cells. The experiments included 5 independent biological samples and 2 technical replicates for each sample for each genotype.

### Chondrocyte metabolic profiling analysis

Passage 1 WT and *Sirt5*^*−/−*^ mouse primary chondrocytes were cultured in 6 cm Petri dishes to full confluence. Cells were then treated with or without 20 mM glucose and 20 nM insulin for 24 hours prior to collection for Gas/Chromatography-Mass/Spectrometry (GC-MS) semi-targeted metabolomics analysis (23). Cells were collected by quickly aspirating the medium, washing twice with warm (37 °C) ultra-pure water to remove medium residuals, immediately snap-freezing with liquid nitrogen, and storing frozen cell samples at −80 °C until extraction and analysis. Metabolite extraction, derivatization, and GC-MS analysis were performed as previously described (23). The analysis also included sample blanks, three pooled quality controls (QC), and external standards of pyruvate, lactate, succinate, fumarate and glucose (Sigma-Aldrich, St. Louis, USA). Metabolites were annotated using NIST library integrated to Agilent MassHunter software, the aforementioned external standards, and our previously made in-house library. Relative abundance of annotated metabolites was calculated by peak area and normalized by internal standards (ribitol).

### Statistical analysis

Semi-quantitative histological scoring data for SIRT5 staining were shown as mean ± 95% confidence interval, and were statistically evaluated using a non-parametric method for two sample comparisons (Mann-Whitney). Semi-quantitative histological scoring data for MaK staining were plotted as mean ± SD and were statistically evaluated using a non-parametric method for two sample comparisons (Mann-Whitney). Data for cellular MaK and COLII staining intensity in different metabolic conditions were plotted as mean ± SD and statistically evaluated using 1-way ANOVA followed by Tukey’s multiple comparison test with a single pooled variance. Parameters in Glycolytic Rate Assay and Mito Stress Assay were plotted as mean ± SEM, and evaluated by unpaired Student’s t-test. Principal component analysis (PCA) was performed on Log-transformed data using a built-in “stat”-package under the R-environment (https://www.r-project.org). Statistically significant changes in Log-transformed metabolite levels were determined by the Student’s t-test (p<0.05, n=3 per group), and data were visualized by heat-map using “ggplot2” package (https://cran.r-project.org/web/packages/ggplot2/citation.html) under the R-environment. Pathway analysis was performed using MetaboAnalyst online platform (https://www.metaboanalyst.ca) and KEGG database (https://www.genome.jp/kegg/pathway.html).

## RESULTS

### Lysine malonylation (MaK) is upregulated in cartilage of db/db mice

To examine if obesity alters lysine malonylation (MaK) in cartilage tissue, we conducted immuno-staining for MaK in knee articular cartilage from obese *db/db* mutant mice and age-matched controls. We observed robustly elevated MaK staining in the chondrocytes from the tibia and femur of *db/db* mice (Figure 1A). Notably, the majority of positively stained chondrocytes were limited to non-calcified cartilage with little staining found in calcified cartilage located below the tidemark (Figure 1A). Within the non-calcified cartilage, MaK staining tended to be more intense in chondrocytes residing in the middle-deep zones. MaK staining was minimal in WT animals, resembling staining observed in isotype controls (Figure 1B). We used a semi-quantitative scoring method to compare the overall staining differences. The average score of MaK staining intensity in tibia of *db/db* mice was 1.75 out of a maximum score of 2.0, which was more than 9 times higher than that in the same compartment of WT mice (Figure 2C). Quantification of MaK staining intensity in femur showed a similar upregulation in *db/db* mice, although the difference was smaller.

**Figure 1.**
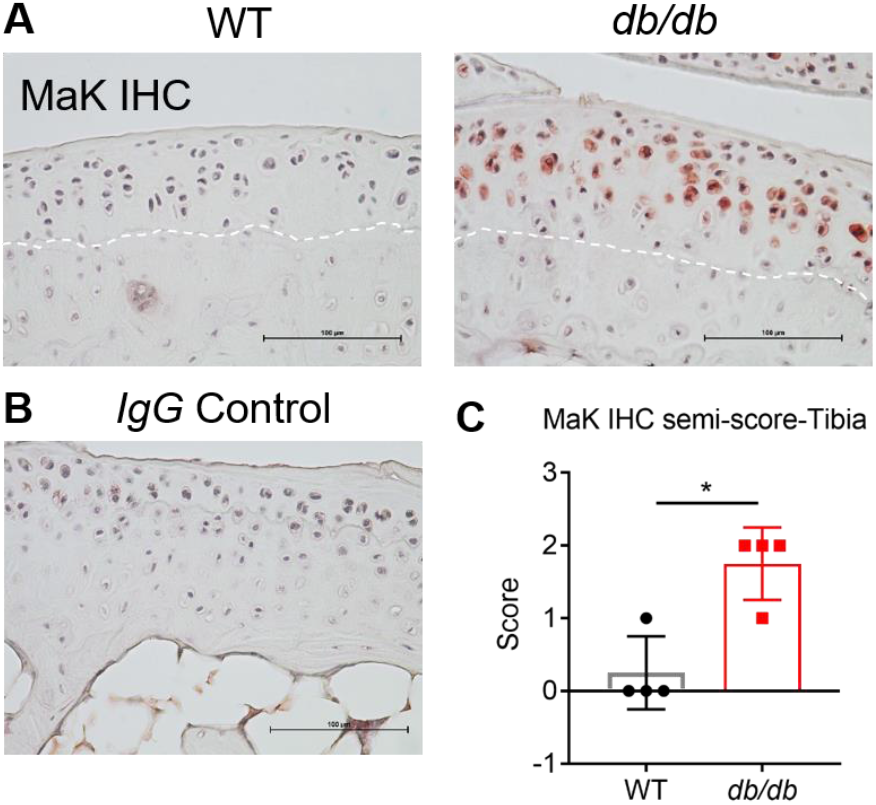
Lysine malonylation (MaK) is upregulated in cartilage of *db/db* mice. **(A)** Representative immunohistochemical staining for malonyl lysine (MaK) in proximal tibial cartilage of 24-week-old male WT (left) and leptin receptor mutant *db/db* (right) mice shows a robust upregulation of protein lysine malonylation in *db/db* chondrocytes. Dashed lines indicate the tidemark, which separates non-calcified cartilage (above) and calcified cartilage (below). **(B)** IgG negative control using samples from *db/db* mice. **(C)** Semi-quantitative scoring of malonyl lysine staining intensity in proximal tibial cartilage. Data points show average scoring results for 4 independent biological samples. 0=no staining; 1=light staining; 2=strong staining. Bars are mean ± SD. *p ≤ 0.05.

**Figure 2.**
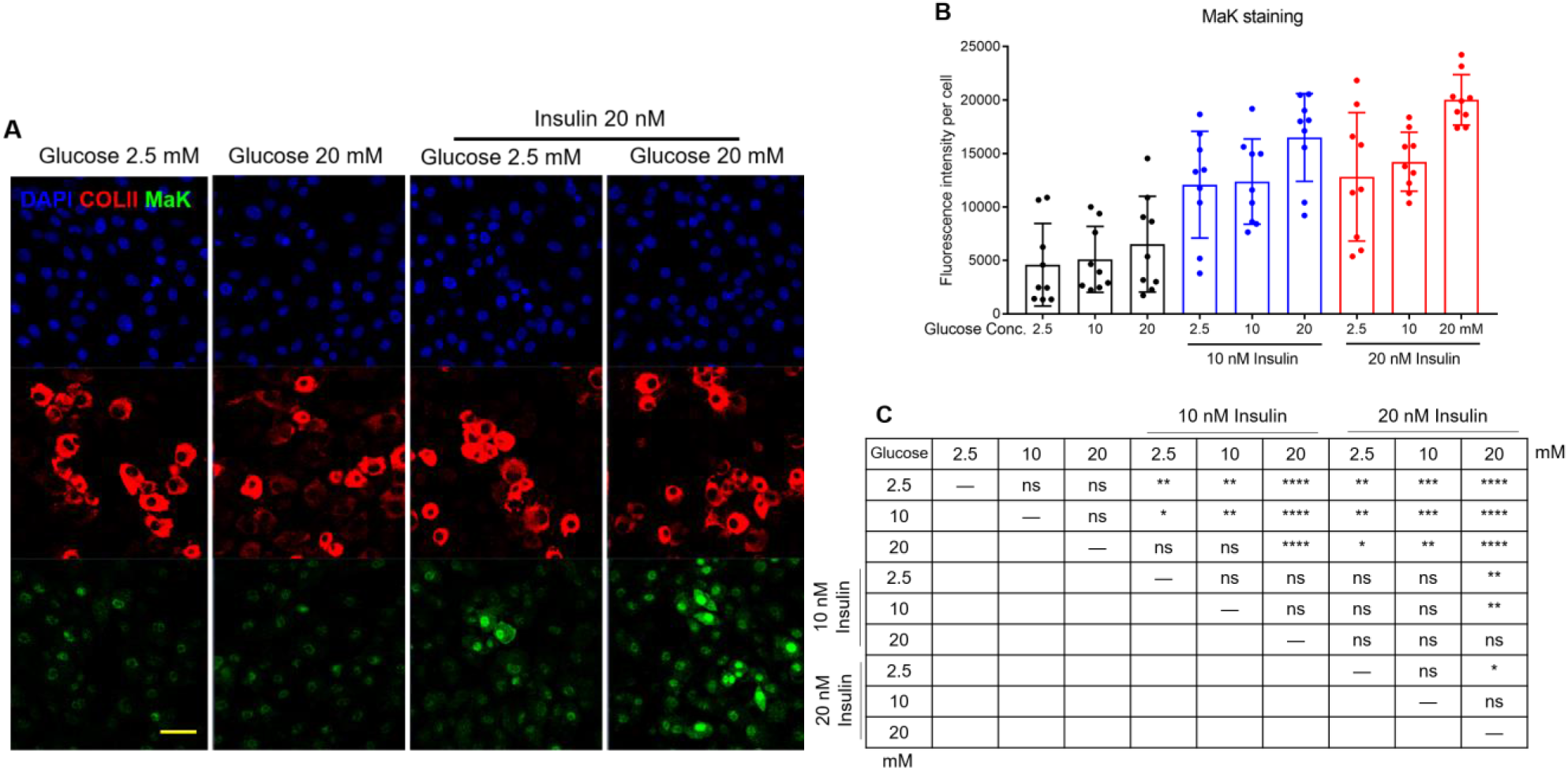
High glucose and high insulin treatment (Glu^hi^Ins^hi^) promotes the accumulation of MaK in cultured chondrocytes. **(A)** Representative image of immunofluorescent staining for MaK (green) and type 2 collagen (COLII, red) in chondrocytes treated with different concentrations of glucose (2.5, 20 mM) with or without insulin (20 nM). **(B)** Quantification of DAPI-normalized MaK staining intensity in chondrocytes treated with different concentrations of glucose (2.5, 10, 20 mM) and insulin (0, 10, 20 nM) (n=9). **(C)** Table summary of statistical analysis. Data shown as mean ± SD. *p≤ 0.05, **p ≤ 0.01, ***p ≤ 0.001, ****p≤ 0.0001.

### High glucose and high insulin treatment (Glu^hi^Ins^hi^) increases MaK level in chondrocytes in vitro

To gain mechanistic information regarding the role of MaK in chondrocyte metabolism, we modelled obese-related metabolic conditions *in vitro*. We challenged primary chondrocytes with different concentrations of glucose and insulin. Cells were then immunostained to detect changes in cellular MaK. Based on cell-normalized MaK immunofluorescence intensity, we found that high glucose treatment alone did not substantially alter chondrocyte MaK levels; whereas, insulin at either 10 or 20 nM significantly increased MaK (Figure 2A, 2B). Interestingly, a combination of 20 mM glucose and 20 nM insulin (Glu^hi^Ins^hi^) condition induced the highest level of MaK in chondrocytes (Figure 2B). The 10 nM insulin with 20 mM glucose and 20 nM insulin with 10 mM glucose also induced MaK levels similar to the Glu^hi^Ins^hi^ treatment condition (Figure 2B, 2C). Glucose and insulin treatment did not alter the expression of COLII, a marker of mature chondrocytes (Figure 2A and Supplementary Figure 1).

### Aging and joint instability-induced OA alter SIRT5 cartilage expression in vivo

The level of MaK in cells is regulated by the non-enzymatic reaction of malonyl-CoA with reactive protein lysines and its reversible removal by Sirt5. Sirt5 is the major demalonylase enzyme, and deficiency in Sirt5 causes hypermalonylation of proteins that are enriched in glycolytic pathway (17). To examine the potential dysregulation of the Sirt5-MaK metabolic pathway in cartilage under OA risk conditions, we performed Sirt5 immuno-staining in cartilage from 22- and 56-week-old mice 8 weeks after knee DMM surgery. Results were compared across age and with the age-matched non-surgical contralateral knee. Chondrocyte SIRT5 staining was detected in nearly all samples from the medial (Figure 3A) and lateral (Figure 3B) knee compartments. The positive staining occurred throughout the cellular space of chondrocytes, consistent with ubiquitous expression of SIRT5 in different cellular compartments including cytoplasm, mitochondria, and nucleus. In the medial compartment, semi-quantitative scoring of combined femoral and tibial cartilage staining showed a trend for reduced Sirt5 staining with increased age in the non-surgical control knee (p=0.06; Figure 3C). However, in the DMM knee, Sirt5 immunostaining did not decrease with age. Consequently, in the 56-week-old animals, Sirt5 staining was elevated in the DMM knee relative to the non-surgical control. The lateral joint compartment, which develops minimal OA changes following DMM surgery, did not show these age-related changes in Sirt5 staining intensity (Figure 3C). To further evaluate the potential involvement of Sirt5 in OA development, we checked for OA-related Gene Expression Omnibus (GEO) datasets in the National Center for Biotechnology Information website. We identified a dataset uploaded by Dr. Frank Beier and colleagues (24). The experimental design compared global gene expression between normal and OA cartilage in rats following an anterior cruciate ligament transection (ACLT) and partial medial meniscectomy. These data showed that Sirt5 mRNA expression was significantly downregulated in OA cartilage compared to either sham or contra-lateral control cartilage (Figure 3F).

**Figure 3.**
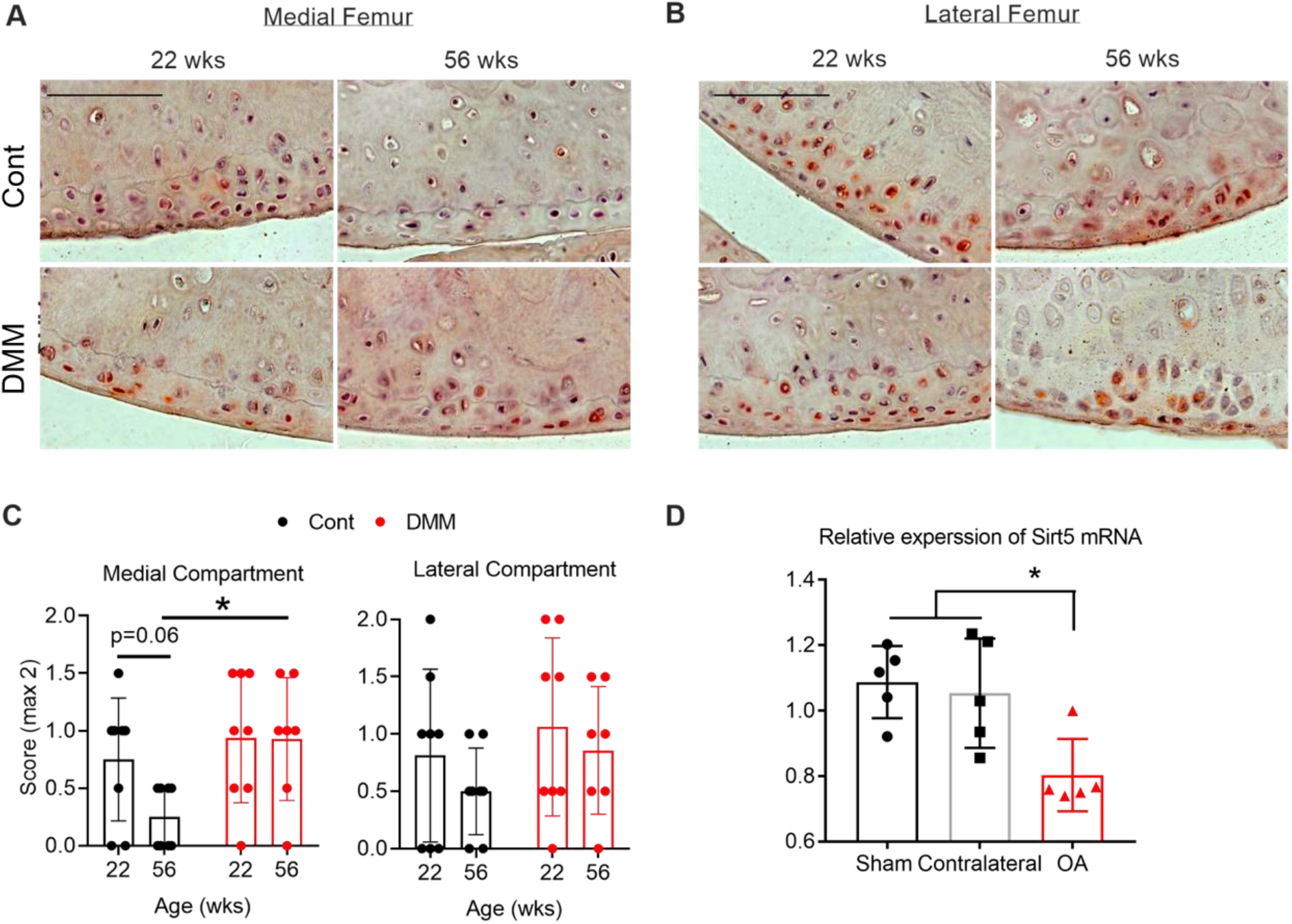
Effect of aging and joint instability-induced OA on cartilage SIRT5 expression. Representative images of immunohistochemical staining for SIRT5 in articular cartilage from (**A**) the medial femur and (**B**) the lateral femur of 22- and 56-week-old mice 8 weeks after DMM surgery. The contra-lateral knee served as the non-surgical control (Cont). Scale bar=50 μm. **(C)** Semi-quantitative scoring of SIRT5 staining intensity in chondrocytes from the femoral and tibial articular cartilage, shown separately for medial and lateral joint compartments. Data points are the average scores of 8 animals per age. 0=no staining; 1=light staining; 2=strong staining. **(D)** Relative Sirt5 expression plotted from Gene Expression Omnibus (GEO) datasets uploaded by *Appleton and colleagues, Arthritis & Rheumatology, 2007* (24). These data were obtained from the cartilage of male Sprague-Dawley rats (350g) that were subject to ACL transection and partial medial meniscectomy OA surgery or Sham surgery. 4 weeks after surgery, cartilage RNA was collected from OA joints, contralateral joints, and joints from Sham group and used for whole genome array. Unpaired t tests were conducted Sham vs OA and Contralateral vs OA. *p<0.05, n=5 per group.

### Increased protein malonylation in cartilage of mice lacking Sirt5

Although Sirt5 has been shown to function as a demalonylase in liver, the role of Sirt5 as a demalynolase enzyme in other tissues, including cartilage, is not well understood. To determine if Sirt5 is also a demalonylase in chondrocytes, we used western blotting to probe the MaK level in WT and *Sirt5*^*−/−*^ chondrocytes isolated from Sirt5 global knockout mice. Compared with WT chondrocytes, there is a significant increase in MaK from total protein homogenates of *Sirt5*^*−/−*^ chondrocytes (Figure 4A). In agreement with previous findings, the differentially malonylated proteins span a wide range of molecular weights (Figure 4A), suggesting that Sirt5 deficiency resulted in global protein hyper-malonylation in chondrocytes. Quantification of the overall intensity of MaK normalized to actin showed that the MaK level was approximately doubled in *Sirt5*^*−/−*^ chondrocytes compared with WT chondrocytes (Figure 4B). Notably, the level of MaK in *Sirt5*^*−/−*^ chondrocytes was more variable among the independent biological replicates compared to WT chondrocytes, suggesting that multiple factors contribute to MaK homeostasis.

**Figure 4.**
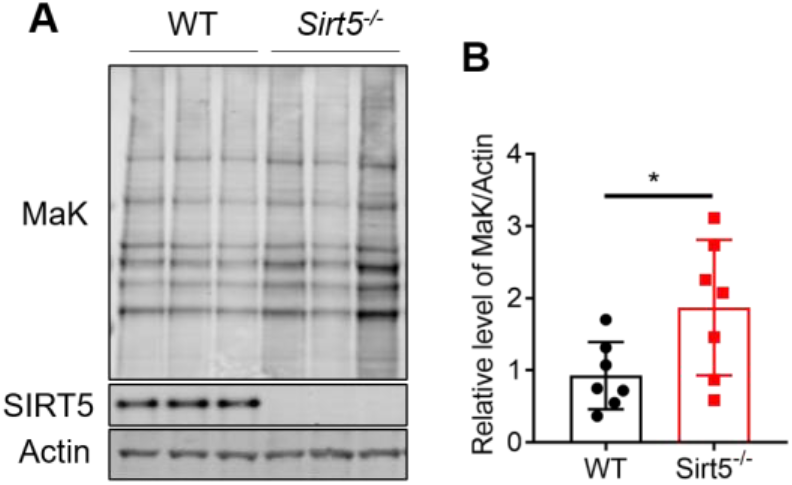
Sirt5 deficiency leads to increased MaK in chondrocytes. **(A)** Representative gel of Western Blotting analysis of MaK, SIRT5, and Actin in chondrocytes isolated from *Sirt5*^*−/−*^ or WT mice. **(B)** Quantitative results of densitometry analysis of MaK in chondrocytes normalized to actin. *p=0.035, primary cells derived from n=7 animals per genotype.

### Sirt5 deficiency reduces chondrocyte glycolysis

Chondrocytes rely on glycolysis for most cellular ATP production. Considering the previously identified effect of Sirt5 deficiency on increased MaK of glycolytic enzymes in the liver and reduced glycolytic flux rate in *Sirt5*^*−/−*^ primary hepatocytes (17), we wanted to evaluate if Sirt5 deficiency also impaired glycolytic metabolism in primary chondrocytes. Therefore, we used a Seahorse XFe24 Analyzer to measure extracellular acidification rate (ECAR) in primary WT and *Sirt5*^*−/−*^ chondrocytes. The ECAR assay measures the concentration of dissolved free protons generated largely by glycolysis as an indirect measure of the overall glycolytic rate. As in hepatocytes, ECAR was reduced in *Sirt5*^*−/−*^ chondrocytes compared to cells from WT animals (Figure 5A). The basal glycolytic rate was nearly 20% lower in *Sirt5*^*−/−*^ chondrocytes compared to WT chondrocytes (Figure 5B). The compensatory glycolysis rate was evaluated following the addition of Rotenone and Antimycin A to inhibit mitochondrial electron transport chain function, forcing a compensatory use of glycolysis to meet cellular energy demands. This compensatory response was also reduced in *Sirt5*^*−/−*^ chondrocytes (Figure 5C). Following rotenone and antimycin A treatment, ECAR transiently increased by 10.4 pmol/min/μg in *Sirt5*^*−/−*^ chondrocytes compared to 18.6 pmol/min/μg in WT chondrocytes (p=0.016; Supplemental Figure 2C). The extracellular acidification can potentially come from 2 sources: conversion of uncharged glucose or glycogen to lactate^−^ + H+ and production of CO_2_ during substrate oxidation and further hydration to HCO3^−^ + H^+^. We checked whether the differences in ECAR included differences in respiration-associated carbonic acid production by measuring the glycolysis-dependent proton efflux rate. We found that nearly all of the proton efflux rate in chondrocytes was attributed to glycolysis and did not differ between WT and *Sirt5*^*−/−*^ chondrocytes (percent proton efflux from glycolysis: WT = 95.8%, *Sirt5*^*−/−*^ = 94.8%; Supplemental Figure 2A). Interestingly, we also observed a lower rate of post 2D-glucose acidification in *Sirt5*^*−/−*^ chondrocytes (Supplemental Figure 2B), which accounted for a third of the difference in basal glycolysis between WT and *Sirt5*^*−/−*^ chondrocytes.

**Figure 5.**
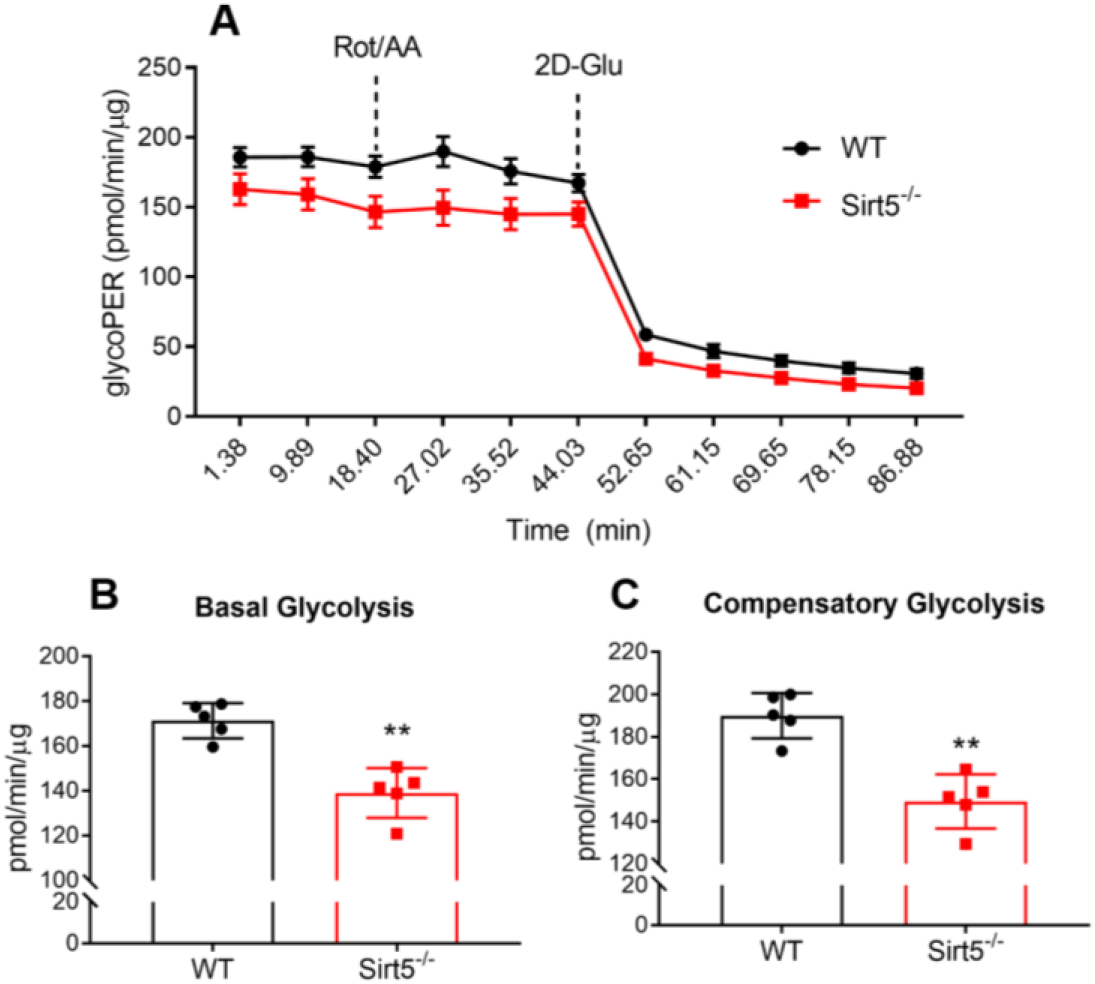
*Sirt5*^*−/−*^ chondrocytes have lower glycolytic rate. **(A)** Extracellular acidification rate as a function of time of WT and *Sirt5*^*−/−*^ chondrocytes as measured by Seahorse respirometry analyzer (data normalized to total protein content). Calculated individual parameters: (**B**) Basal Glycolysis and (**C**) Compensatory Glycolysis. Data shown as mean ± SD. **p<0.01, n=5 independent biological replicates per genotype.

### Sirt5 deficiency reduces chondrocyte basal and ATP-linked cellular respiration

Sirt5 may also translocate to the mitochondria (25) and regulate mitochondrial PTMs (17). Thus, to comprehensively understand the effect of Sirt5 deficiency on chondrocyte cellular metabolism, we used Seahorse Respirometry to assess changes in chondrocyte cellular respiration. With sequential addition of the ATP synthase inhibitor oligomycin, protonophoric uncoupler FCCP, and electron transport inhibitors rotenone and antimycin A, it is possible to test for major differences in mitochondrial function between WT and *Sirt5*^*−/−*^ chondrocytes by measuring the oxygen consumption rate (OCR) (Figure 6A). Before oligomycin was added, OCR was 20% lower in *Sirt5*^*−/−*^ chondrocytes compared to WT chondrocytes (Figure 6B). To understand if this difference was linked to ATP production, we compared the difference in OCR before and after oligomycin was added. We found that WT and *Sirt5*^*−/−*^ chondrocytes had similar rates of respiration after the addition of oligomycin (Figure 6C), suggesting that the defect of basal respiration in *Sirt5*^*−/−*^ chondrocytes is due to decreased ATP production. ATP synthase inhibition by oligomycin will significantly reduce electron flow through ETC, however electron flow is not fully stopped due to a process known as proton leak. We estimated proton leak based on OCR following ATP synthase inhibition. Less than 10% of OCR was estimated to be from proton leak in both *Sirt5*^*−/−*^ and WT chondrocytes, and this value was unaltered by Sirt5 deficiency (Supplemental Figure 3A). This observation was also reflected by a similar coupling efficiency, calculated by the percentage of ATP production rate to basal respiration rate, between *Sirt5*^*−/−*^ and WT chondrocytes (Figure 6E). Additionally, analysis of OCR after the uncoupler FCCP was added showed that the maximal respiration rate (Figure 6D) as well as spare respiratory capacity comparing (Supplemental Figure 3B) were lower in *Sirt5*^*−/−*^ chondrocytes compared with WT chondrocytes. Together, these results indicate that Sirt5 deficiency in chondrocytes also alters mitochondrial function.

**Figure 6.**
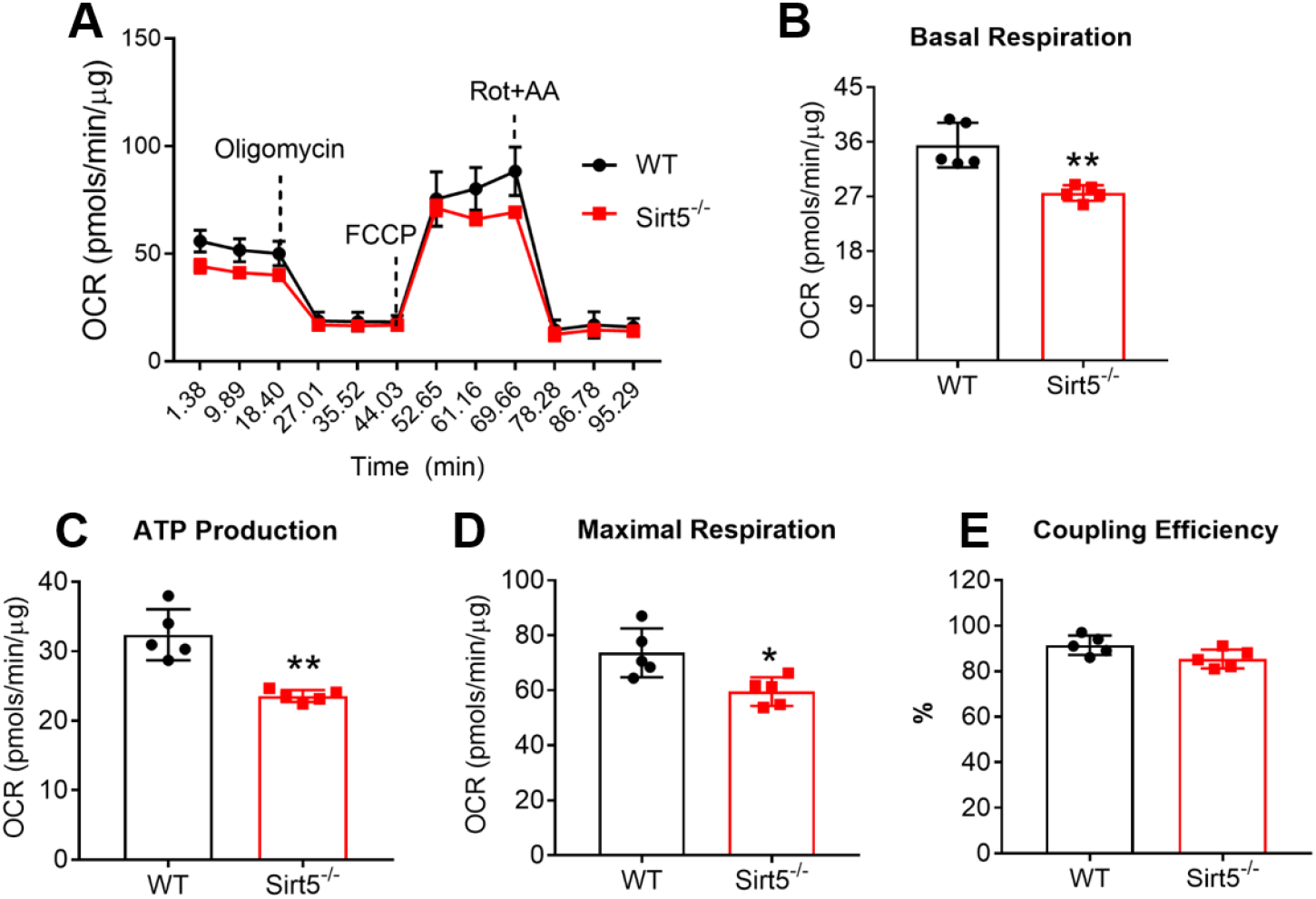
*Sirt5*^*−/−*^ chondrocytes have lower oxygen consumption rate associated with basal respiration and ATP production. **(A)** Oxygen consumption rate as a function of time of WT and *Sirt5*^*−/−*^ chondrocytes measured by Seahorse respirometry analyzer (data normalized to total protein content). Calculated individual parameters: (**B**) Basal Respiration, (**C**) ATP Production, (**D**) Maximal Respiration, and (**E**) Coupling Efficiency. Data shown as mean ± SD. **p<0.01, *p<0.05, n=5 independent biological replicates per genotype.

### Metabolomics analysis reveals distinct cellular metabolism in *Sirt5*^*−/−*^ and WT chondrocytes

Considering Sirt5 has been identified to regulate PTMs of a wide range of proteins across different metabolic pathways, we conducted a high-throughput GC-MS analysis of small molecules (metabolites) to more broadly evaluate the role of Sirt5 in chondrocyte cellular metabolism. The experimental groups were designed to evaluate the qualitative and quantitative changes of metabolites in WT and *Sirt5*^*−/−*^ chondrocytes treated with or without Glu^hi^Ins^hi^. We included three biological replicates in each group.

A total of 41 primary metabolites were detected and identified (see Methods section) across all four groups. Principle component analysis (PCA) of the results demonstrated a clear and strong separation of non-treated and treated groups by the 1^st^ principal component (Figure 7A, PC1, ~84% of variance explained), suggesting a strong effect of Glu^hi^Ins^hi^ treatment on chondrocyte cellular metabolism. Additionally, PCA analysis revealed a clear separation between WT and *Sirt5*^*−/−*^ chondrocytes through the 2^nd^ Principal component (Figure 7A, PC2, ~9% of variance explained), suggesting that *Sirt5*^*−/−*^ and WT chondrocytes are metabolically distinct. Notably, the greatest qualitative differences between WT and Sirt5 deficient chondrocyte metabolic profiles occurred with the Glu^hi^Ins^hi^ treatment. This observation was supported by a statistical analysis of the difference in abundance of individual metabolites between WT and *Sirt5*^*−/−*^ chondrocytes. 13 metabolites were significantly different under basal conditions while 18 were different following Glu^hi^Ins^hi^ treatment (Supplemental Figure 4). Interestingly, under the basal condition, only 3 out of the 13 metabolites were upregulated in *Sirt5*^*−/−*^ chondrocytes (Supp Figure 4). In contrast, under Glu^hi^Ins^hi^ treatment, 9 of the 18 significantly changed metabolites were upregulated in *Sirt5*^*−/−*^ chondrocytes (Supp Figure 4). This suggests that Sirt5 deficiency may target different metabolic pathways under different metabolic conditions. Indeed, pathway analysis showed that under the basal condition, Sirt5 deletion primarily altered amino acid (including branched-chain amino acids and aromatic amino acids) metabolism and glutathione metabolism (Figure 7B). Whereas under Glu^hi^Ins^hi^ treatment, a wide range of metabolic pathways, including aromatic amino acids, TCA cycle, alanine/aspartate/glutamate metabolism, and arginine biosynthesis, were significantly impacted by Sirt5 deficiency (Figure 7B).

**Figure 7.**
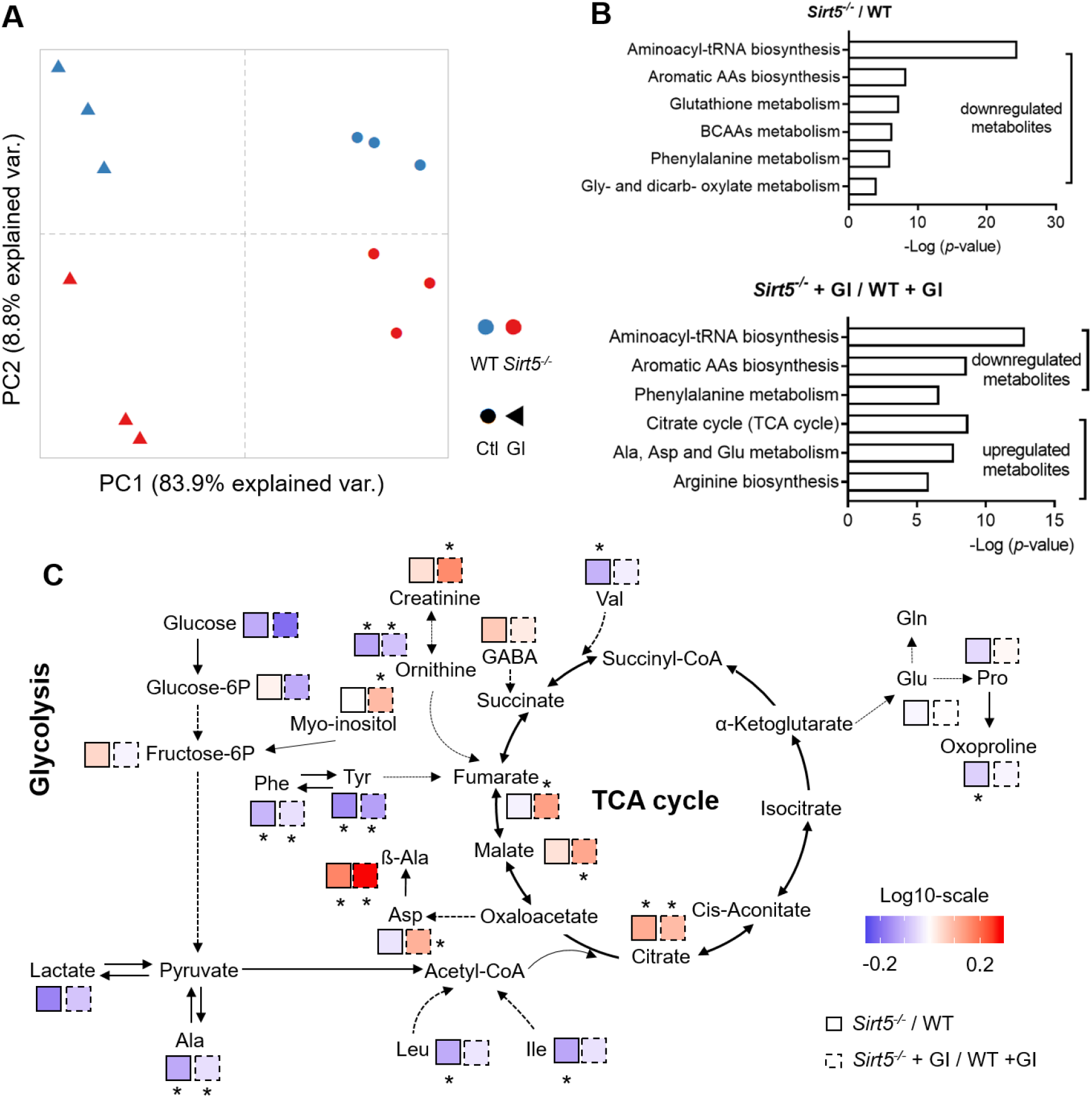
GC-MS analysis of relative content of metabolites in chondrocytes showed an effect of Sirt5 deficiency and Glu^hi^Ins^hi^ condition on chondrocyte cellular metabolism. **(A)** Principal component analysis (PCA) results based on metabolite abundance values for metabolites detected in both WT and Sirt5*-/-* chondrocytes treated with or without Glu^hi^Ins^hi^. (B) Pathway analysis of significantly changed metabolites comparing Sirt5^−/−^ and WT chondrocytes treated with or without Glu^hi^Ins^hi^ (GI), using MetaboAnalyst web-platform (https://www.metaboanalyst.ca) on KEGG database background. **(C)** Metabolic map (based on KEGG database) of the majority of detected metabolites in WT and Sirt5^−/−^ chondrocytes. Significantly changed metabolites in the map were marked as *p<0.05. Icons with solid outlines indicate non-treated groups, icons with dashed outlines indicate GI treated groups.

To better understand the metabolic changes associated with Sirt5 deficiency, we utilized a systematic approach of metabolic profiling through mapping individual metabolites, including those that did not show significant alterations, in the context of a metabolic network. We visualized the abundance of metabolites in this network by heat-map icons indicating relative changes between *Sirt5*^*−/−*^ and WT within each treatment. Overall, the results of this analysis demonstrated contrasting differences among multiple metabolic pathways. Though few metabolites involved in glycolysis were significantly altered by Sirt5 deficiency, several metabolites related to glucose metabolism were downregulated (blue) in *Sirt5*^*−/−*^ chondrocytes regardless of Glu^hi^Ins^hi^ treatment (Figure 7C). By contrast, 3 TCA cycle metabolites (fumarate, malate, and citrate) showed upregulation in *Sirt5*^*−/−*^ (Figure 7C). Another interesting observation was the widespread change in amino acids levels with Sirt5 deletion. More specifically, the branched-chain amino acids leucine, isoleucine and valine were significantly reduced in *Sirt5*^*−/−*^ chondrocytes under basal condition. These changes were diminished when Glu^hi^Ins^hi^ treatment was applied (Figure 7C). In addition, the aromatic amino acids tyrosine and phenylalanine were also reduced in *Sirt5*^*−/−*^ chondrocytes (Figure 7C). These results are consistent with the pathway analysis results indicating that aromatic amino acids biosynthesis and branched-chain amino acids degradation pathways were altered by Sirt5 deletion (Figure 7B).

## Discussion

In this study we describe for the first time that Sirt5 regulates chondrocyte metabolism and has altered expression in OA cartilage tissue, suggesting that Sirt5 and its enzymatic target, post-translational protein lysine-malonylation (MaK), are involved in musculoskeletal disease development. Recent studies have identified post-translational modifications (PTMs) of proteins at lysine residues via reversible acylation (e.g., malonylation, succinylation, and acetylation) as important regulators of enzymatic activity and cellular metabolism (13, 14). These PTMs are regulated by deacylase sirtuins, which can remove the acyl group from proteins and restore their enzymatic activity. Dysregulation of sirtuins and PTMs has been implicated in the pathogenesis of many diseases. In the current study, we hypothesized that Sirt5-mediated MaK regulates chondrocyte cellular metabolism under basal and obesity-associated conditions. We found that protein malonylation was significantly increased in cartilage of obese mice and in primary chondrocytes cultured with high concentrations of glucose and insulin (Glu^hi^Ins^hi^) to model obesity-related conditions. These findings indicate that chondrocytes increase acylation-based PTMs like other metabolically active tissues under obesity conditions. We further found that Sirt5 deficiency in chondrocytes causes hyper-malonylation of global cellular proteins and alters cellular metabolism via changes in a wide range of metabolic pathways, including TCA cycle, glycolysis, and amino acid metabolism.

The 7 sirtuins (Sirt 1-7) found in mammals vary in their enzymatic activities to different forms of PTMs. SIRT5 was initially reported as a lysine deacetylase based on a conserved deacetylase domain. However, it was later established that SIRT5 efficiently removes acidic acyl groups, such as-succinyl, malonyl, and glutaryl, but not the acetyl moiety from lysine residues. Thus, SIRT5 has now been considered as a potent desuccinylase, demalonylase, and deglutarylase (26). We found that Sirt5 deficiency in chondrocytes leads to hypermalonylation of global proteins, suggesting Sirt5 is also a demalonylase in cartilage tissue. We did not evaluate other Sirt5-dependent PTM acyl moieties, such as lysine succinylation or glutarylation, although they could also be mechanisms by which Sirt5 regulates chondrocyte function. For example, SIRT5 has been found to directly desuccinylate multiple TCA cycle enzymes, including succinate dehydrogenase subunit A (SDHA) (27) and isocitrate dehydrogenase 2 (IDH2) (28). Additionally, Sirt5 has also been shown to play a role in regulating glutamine metabolism in colorectal cancer by direct deglutarylation and activation of glutamate dehydrogenase 1 (GLUD1) (29). Further studies on identifying the role of Sirt5 mediated regulation of other types of PTMs in chondrocytes are warranted.

In the current study, we discovered that there was a robust elevation of protein malonylation in the cartilage of *db/db* obese mice. This upregulation can also be recapitulated *in vitro* by culturing primary chondrocytes with high concentrations of glucose and high insulin. The biological significance of lysine malonylation remains poorly understood, due at least in part to a lack of knowledge of the proteins and lysine residues that become modified. Du et al. recently reported the hypermalonylation of proteins in livers of *db/db* diabetic mice. Using affinity enrichment coupled with proteomic analysis, they identified 573 malonylation sites over 268 proteins, which are enriched in glucose and fatty acid metabolism pathways. Interestingly, Sirt5 mediated protein malonlylation was also observed to target glycolysis pathway enzymes in the liver (17). This suggests that conserved targets for Sirt5-regulated lysine malonylation might exist under different metabolic conditions. It is of great interest to understand how Sirt5 regulates desuccinylation and demalonylation of specific proteins in chondrocytes. Proteomics-based screening and identification of metabolic target enzymes in chondrocytes for Sirt5 mediated desuccinylation and demalonylation is an important next step.

A novel finding from this study is that deleting Sirt5 in chondrocytes leads to a significantly lower glycolytic flux rate as well as lower basal respiration rate as measured by Seahorse Respirometry. We found that Sirt5 expression is altered in OA cartilage, suggesting that Sirt5 may contribute to changes in chondrocyte metabolism during the pathogenesis of OA. Indeed, Sirt5 mediated PTMs have been demonstrated to target a variety of important enzymes in different metabolic pathways including glycolysis, TCA cycle, and electron transport chain. For example, SIRT5 was shown to directly demalonylate GAPDH and likely other glycolytic enzymes in mouse liver, leading to increased GAPDH activity and elevated glycolytic flux (17). Moreover, SIRT5 was found to desuccinylate pyruvate kinase (PKM2) in A549 NSCLC and 293T cell lines, facilitating the conversion of phosphoenolpyruvate to pyruvate in the final step of glycolysis (30). These findings are consistent with our observation that Sirt5 regulates both glycolysis and oxidative phosphorylation in chondrocytes.

We found that Sirt5 is abundantly expressed in cartilage and changes in expression following an injury in mouse and rat models of post-traumatic OA. Previous studies have reported changes in other sirtuins during the development of OA. For example, the nuclear deacetylase Sirt1 was reduced at the protein level in human OA chondrocytes (31). Sirt3, a mitochondrial protein deacetylase, was reduced at the protein level in mouse, rat, and human cartilage with increasing age (11). Our studies showed that Sirt5 cartilage protein level, as determined by immunohistochemistry, were increased in medial femoral and tibial cartilage 8 weeks following DMM surgery compared to site-matched cartilage from the contra-lateral control knee. This increase occurred in older mice only, suggesting an important age-dependent component in Sirt5 changes with OA. Interestingly, using publicly available data (24), we observed a reduction in Sirt5 gene expression in a rat model of post-traumatic OA (anterior cruciate ligament transection and partial medial meniscectomy) 4 weeks after injury and forced mobilization compared to cartilage from the contra-lateral knee or sham surgery. These differences in expression may be due to many factors, such as species, age, injury model, and timepoint of analysis post injury. In the future, studies including both gene and protein expression in the same animal model at multiple time points will help to resolve the current uncertainty in how Sirt5 changes during the course of OA progression.

One way to examine the biological significance of metabolic proteins PTMs is to measure cellular intermediary metabolites using metabolomics. Metabolic profiling is one of the most used approaches of metabolomics to give an instantaneous snapshot of the cell physiological status and provide a direct functional readout of the underlying biochemical activities of cells/tissues (32). In the current study, the metabolomics analysis revealed a wide range of changed metabolites with Sirt5 deficiency and Glu^hi^Ins^hi^ treatment. Though few glycolytic metabolites significantly changed, many showed a trend for being reduced in *Sirt5*^*−/−*^ chondrocytes. These trends are consistent with previous findings that Sirt5 regulates malonylation and protein-specific activities of glycolytic enzymes (17). Moreover, several detected TCA cycle metabolites, including fumarate, malate, and citrate, were significantly upregulated in Sirt5^*−/−*^ chondrocytes with or without Glu^hi^Ins^hi^ treatment. Additional work is needed to identify the net effect of different forms of Sirt5-dependent PTMs on TCA Cycle enzyme function in chondrocytes, such as SDHA (27) and IDH2 (28). An additional consideration is that changes in the abundance of TCA cycle intermediates could be due to anaplerotic amino acid influx. Sadhukhan et al. discovered that branched-chain amino acid metabolism is a top hit for pathway enrichment of lysine succinylated proteins (33). Additionally, previous studies demonstrated a clear link between branched-chain amino acids and glycolysis regulation (34, 35). These discoveries are consistent with our finding showing that branched-chain amino acid metabolism is significantly downregulated by Sirt5 deficiency under the basal conditions but not under the Glu^hi^Ins^hi^ treatment condition. The semi-targeted metabolomics analysis used in this study identified a limited number of metabolites under steady-state conditions. Future metabolic profiling using a higher-resolution technique or with isotopically-labeled substrates for flux-based approaches would provide additional insight into Sirt5-dependent effects on chondrocyte metabolism.

Although our study focused primarily on Sirt5-dependent demalonylation, our findings raise an important question about how MaK is generated in chondrocytes. From what is known about other PTMs, malonyl-Coenzyme A (malonyl-CoA) likely serves as the universal donor for malonylation reactions either via the activity of a currently unidentified malonyltransferase or via a non-enzymatic mechanism. Cytosolic malonyl-CoA is produced from acetyl-CoA by acetyl-CoA carboxylase 1 (ACC1) when acetyl-CoA exceeds energetic demand (36). Malonyl-CoA produced in the cytosol serves as the building blocks for *de novo* fatty acid synthesis. Meanwhile malonyl-CoA produced by the mitochondrial isoform ACC2 allosterically inhibits carnitine palmitoyl-transferase 1 (CPT1), the rate-limiting mitochondrial fatty acid transporter, and thereby suppresses β-oxidation. The intriguing link between protein malonylation and fatty acid metabolism by the intermediary metabolite malonyl-CoA is relevant to cartilage biology based on the strong association between abnormal metabolic conditions (e.g., obesity and diabetes) and OA (3, 4, 37). Increasing lipid storage has been reported to be positively correlated with OA severity (38) and altered lipid metabolism has been implicated in the pathogenesis of OA (39). Our results showed that high concentrations of glucose and insulin treatment increased protein malonylation. This is consistent with the finding that insulin substantially activates ACC (40) while glucose oxidation provides acetyl-CoA as a substrate for ACC.

We note that our study has two limitations that should be considered. First, the effect of Sirt5-MaK on chondrocyte cellular metabolism was investigated in primary juvenile chondrocytes under *in vitro* cell culture conditions, which differs from *in vivo* physiological environment. We recently reported on the differences between *in vivo* and *in vitro* conditions on cartilage metabolism, which showed enhanced amino acid metabolites under *in vitro* conditions (23). Thus, future *in vivo* strategies are needed to evaluate the broad metabolic effects of Sirt5 deficiency on cartilage. We are currently working to conduct these studies in the joints of *Sirt5*^*−/−*^ mice. An additional limitation of our study was the use of convenience histological samples from prior cohort animal studies to conduct immunostaining analyses (e.g., MaK in Figure 1, SIRT5 in Figure 3). Thus, these analyses were not statistically powered for these outcomes and are discovery-based, requiring future validation.

In summary, this is the first study to evaluate the role of Sirt5 in chondrocyte metabolism. The results extend our understanding of PTMs and sirtuins in regulating chondrocyte function by showing altered levels of Sirt5 and MaK in cartilage with obesity and OA. Our results also provide *in vitro* evidence that modulation of Sirt5-MaK signaling impairs chondrocyte cellular metabolism. These findings provide a new potential mechanism that might be involved in obesity-associated OA development.

## Abbreviations

MaK: lysine malonylation
DMM: destabilization of the medial meniscus
GC-MS: gas/chromatography-mass/spectrometry
Glu^hi^Ins^hi^: high concentrations of glucose and insulin
MetS: Metabolic syndrome
PTM: post-translational modification
GEO: Gene Expression Omnibus
ACLT: anterior cruciate ligament transection
ECAR: extracellular acidification rate
OCR: oxygen consumption rate
PCA: principle component analysis
SDHA: succinate dehydrogenase subunit A
IDH2: isocitrate dehydrogenase 2
GLUD1: glutamate dehydrogenase 1
ACC1: acetyl-CoA carboxylase 1

## Acknowledgement

The authors would like to thank and acknowledge Dr. Erika Lopes and Joanna Hudson for conducting the DMM animal study and tissue collection, Wan-Pin Chang and Joanna Hudson for collecting tissues from the *db/db* mouse study, and Taylor Block for her help with care of Sirt5 knockout mice. The authors also thank Dr. Satoshi Matsuzaki for his help with Seahorse Respirometry analysis. The authors would like to thank the OMRF imaging core facility for their excellent help with confocal imaging.

## Author Contributions

SZ, TMG, KMH, and HVR: Conceptualization; SZ, AB, AJ, and DM: Data acquisition; SZ, AB, and TMG: Data analysis; SZ and AB: Writing-original draft; SZ, AB, AJ, DM, KMH, HVR, and TMG: Writing-review and editing and final approval.

## Supplemental Figures

**Supp Figure 1.**
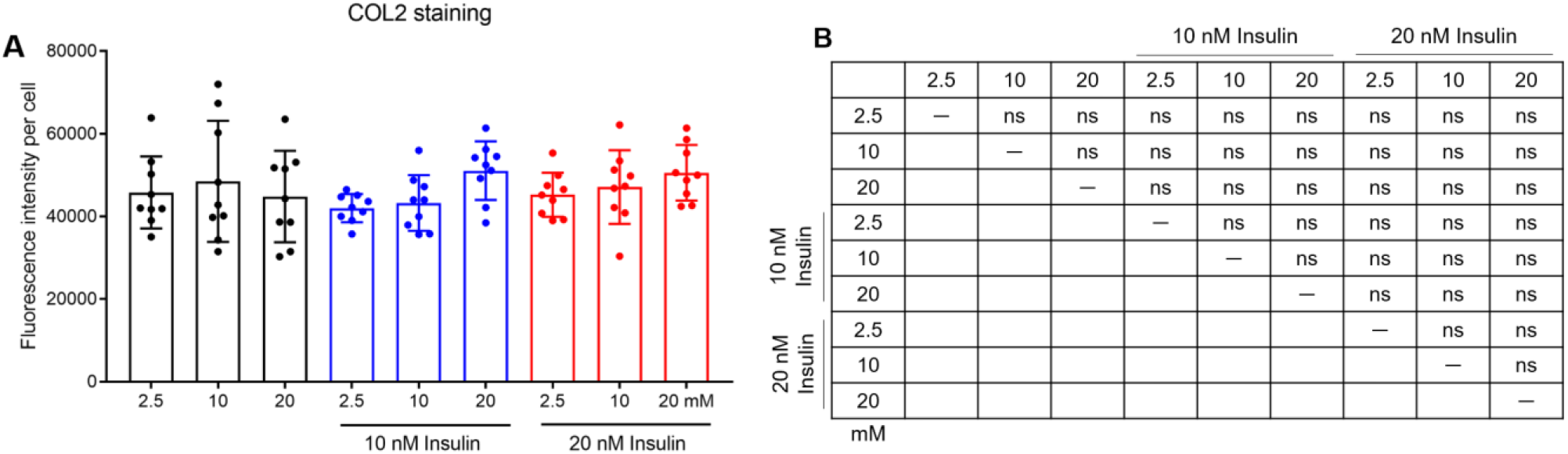
24 hours of high glucose and high insulin treatment (Glu^hi^Ins^hi^) does not alter COL2 protein immunostaining in chondrocytes. **(A)** Quantification of COL2 staining intensity per chondrocyte treated with different concentrations of glucose (2.5, 10, 20 mM) and insulin (0, 10, 20 nM) (n=9). **(B)** Table summary of statistical analysis. Data shown as *mean ± SD.* ns=not statistic significant.

**Supp Figure 2.**
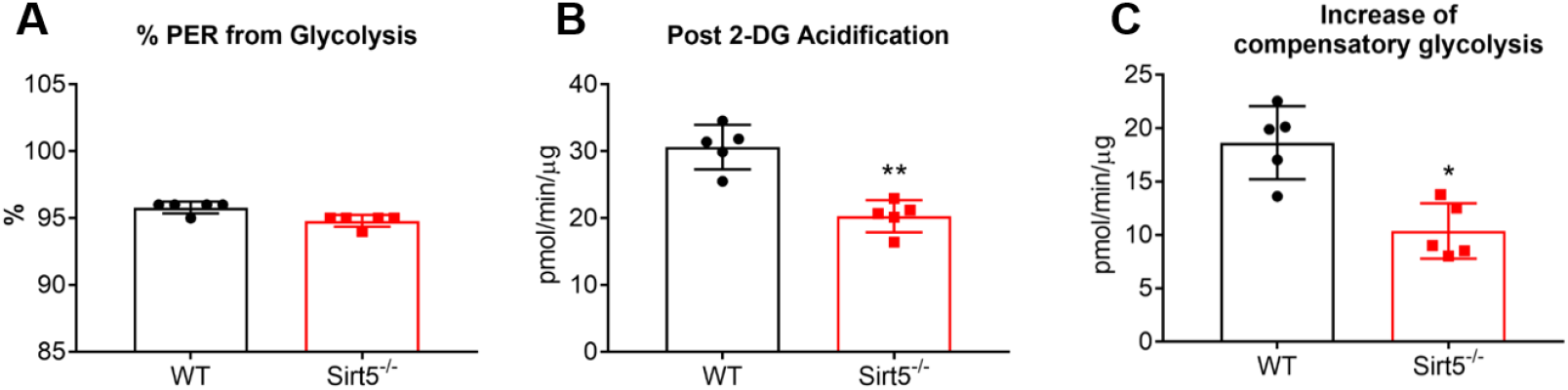
Calculated individual parameters from glycolytic rate Seahorse assay: % PER from glycolysis (A), Post 2-DG acidification (B), and Increase of compensatory glycolysis (C) in *Sirt5*^*−/−*^ versus WT chondrocytes. Data shown as mean ± SD. **p<0.01, *p<0.05, n=5.

**Supp Figure 3.**
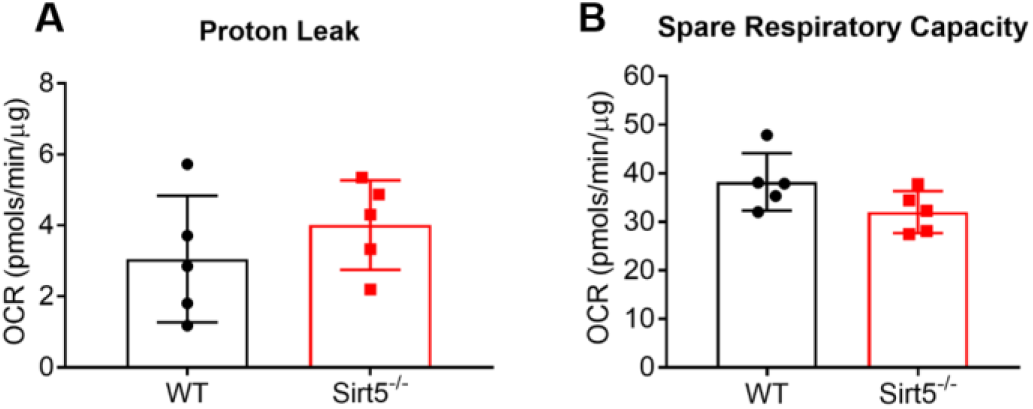
Calculated individual parameters from mito-stress Seahorse assay: Proton leak (A) and Spare Respiratory Capacity (B). Data shown as mean ± SD. n=5.

**Supp Figure 4.**
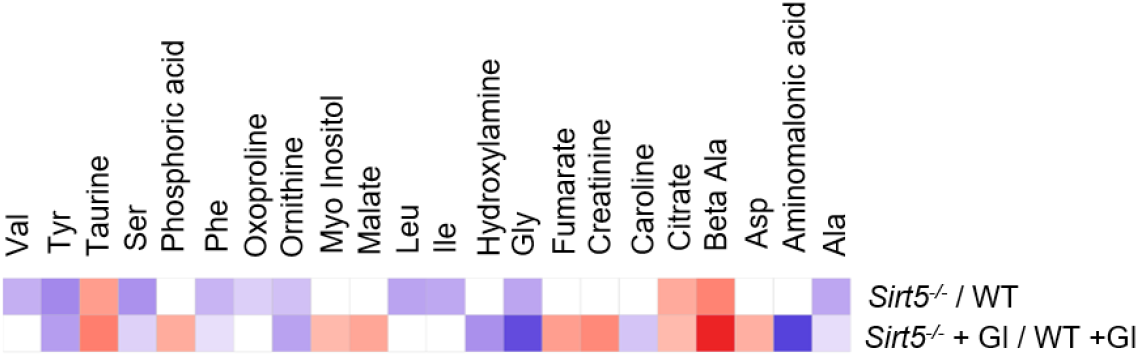
The complete list of significantly changed metabolites in Sirt5^−/−^versus WT chondrocytes treated with or without Glu^hi^Ins^h^ (GI). Only metabolites with colors (blue or red) are significantly changed (p<0.05); metabolites with white color are not significantly changed. Red = upregulated, blue = downregulated.

